# Spatial context non-uniformly modulates inter-laminar communication in the primary visual cortex

**DOI:** 10.1101/2024.02.21.581417

**Authors:** Xize Xu, Mitchell P. Morton, Nyomi V. Hudson, Anirvan S. Nandy, Monika P. Jadi

**Author notes:** Equal contribution (Xize Xu, Mitchell P. Morton). Senior authors (Anirvan S. Nandy, Monika P. Jadi).

## Abstract

Our visual experience is a result of the concerted activity of neuronal ensembles in the sensory hierarchy. Yet how the spatial organization of objects influences neural activity in this hierarchy remains poorly understood. We investigate how the inter-laminar interactions in the primary visual cortex (V1) are affected by visual stimuli in isolation or with flanking stimuli at various spatial configurations that are known to cause non-uniform degradation of perception. By employing dimensionality reduction approaches to simultaneous layer-specific population recordings, we establish that cortical layers interact through a structurally stable communication subspace. Spatial configuration of contextual stimuli differentially modulates inter-laminar communication efficacy, the balance between feedforward and feedback signaling, and contextual signaling in the superficial layers. Remarkably, these modulations mirror the spatially non-uniform aspects of perceptual degradation. Our results suggest a model of retinotopically non-uniform cortical connectivity in the output layers of V1 that influences communication in the sensory hierarchy.

## INTRODUCTION

Spatial vision is the ability to perceive visual objects within three-dimensional space and its dysfunction is detrimental to our ability to interact with the visual world. Our visual experience relies on the coordinated activity of neuronal ensembles in the sensory hierarchy of the cortex (1). Yet how the spatial organization of objects influences interactions between neuronal populations in this hierarchy remains incompletely understood.

Visual perceptual performance has been shown to vary as a function of visual field location, which is best at the center of gaze, degrades with eccentricity, and varies with radial angle. This asymmetry is paralleled by the asymmetric neural organization at multiple stages of the visual system (see (2) for a review). Thus, a comprehensive characterization of the neural correlates and the underlying computations of spatial vision requires empirical investigations without the assumption of spatial isotropy. Phenomena such as visual crowding, the inability to recognize objects amongst clutter, offer a powerful framework for such investigations. Visual crowding is thought to be the primary limitation on object perception in peripheral vision (3), and suggests spatially non-uniform context integration in psychophysical studies (4, 5).

Understanding the neural basis of spatially non-uniform context integration requires identification of where the effects arise in the visual hierarchy, and how information flow along this hierarchy is modulated by context. Neuronal spiking activity recorded from anesthetized monkeys indicates that visual crowding impairs feature representations as early as area V1 (6, 7). Human imaging studies show modulation of activity in area V1 (8, 9), as well as in higher visual areas (10–13). More importantly, inter-areal correlations are disrupted by spatial context integration (14), suggesting modulation of information flow along the hierarchy. Despite extensive psychophysical studies and computational modeling (15–26), the neural basis of the perceptual asymmetry of spatial context integration remains unknown. Understanding how the organization of objects influences information flow along the visual hierarchy is thus critical to our understanding of spatial vision.

Information flow in the visual hierarchy is both inter-areal as well as inter-laminar, since the laminar organization of neuronal populations (1, 27) with their stereotypical patterns of intra- and inter-areal projections (28, 29) is a canonical motif of cortical organization. The neurophysiological aspects of this lamina-specific circuit have been extensively characterized in the context of surround modulation, namely the change in neuronal activity in response to visual stimulation of classical and extra-classical receptive fields (30–39). Geniculate feedforward connections, intra-V1 horizontal connections, and interareal feedback connections to area V1 have all been shown to contribute to surround modulation in V1 at different spatiotemporal scales (see (40) for a review). However, a prevailing assumption in most of these studies has been the spatially uniform nature of the surrounding context. In this study, we investigated how the spatial configuration of visual stimuli modulates inter-laminar interactions in area V1. We specifically focused on interactions between two populations: input layer neurons that receive geniculate inputs and project locally to superficial layers, and superficial layer neurons that project to higher-order visual areas as well as to local deep layers.

We performed laminar recordings from awake macaque V1 with visual stimuli presented either in isolation or with a flanking stimulus at various locations known to cause non-uniform perceptual impairment in peripheral vision. We characterized signaling from the input to the superficial layer which is a key pathway in the feedforward propagation of sensory information. Using dimensionality reduction techniques to identify communication subspaces (41), we found that the input and superficial layers interact through a structurally stable communication subspace under different visual conditions. Flanking stimuli modulated the efficacy of inter-laminar signaling in a location-specific manner, by changing both the subspace efficacy and the balance between feedforward and feedback signaling. Moreover, our analysis revealed a non-uniform contextual signal in the superficial layers triggered by flankers.

## RESULTS

To study inter-laminar interactions in V1 we simultaneously recorded the activity of neuronal populations in the input layer of V1 (unit count: 27.9 ± 3.2 SEM) and their primary downstream target, the superficial layer of V1 (unit count: 21.9 ± 5.0 SEM) in two awake macaque monkeys (Fig. 1A; 14 sessions from Monkey M, 8 sessions from Monkey D). The recorded neurons consisted of well-isolated single units and multi-unit clusters, which had retinotopically aligned receptive fields (Fig. 1B), implying a high probability of direct interactions. Laminar identity was established using current source density (CSD) analysis (42) (Fig. 1C). Monkeys were trained to fixate on the center of the screen and passively view a target stimulus (100ms stimulus duration, 200-250ms inter-stimulus interval) at the receptive field of the recorded cortical column, either in isolation (probe condition) or with a flanking stimulus (flanked condition) at one of four spatial locations relative to the target stimulus (Fig. 1D1), based on which three flanked conditions were defined (Fig. 1D2). In both the radial-in and radial-out flanked conditions, the flanker was positioned on the radial axis connecting the target stimulus and the fixation point. The flanker was placed inward (between the target stimulus and the fixation point) in the radial-in condition while it was placed outward (past the target stimulus) in the radial-out condition. In the tangential flanked condition, the flanker was positioned on either side of the target stimulus along the tangential axis, defined as the axis orthogonal to the radial axis.

**Figure 1.**
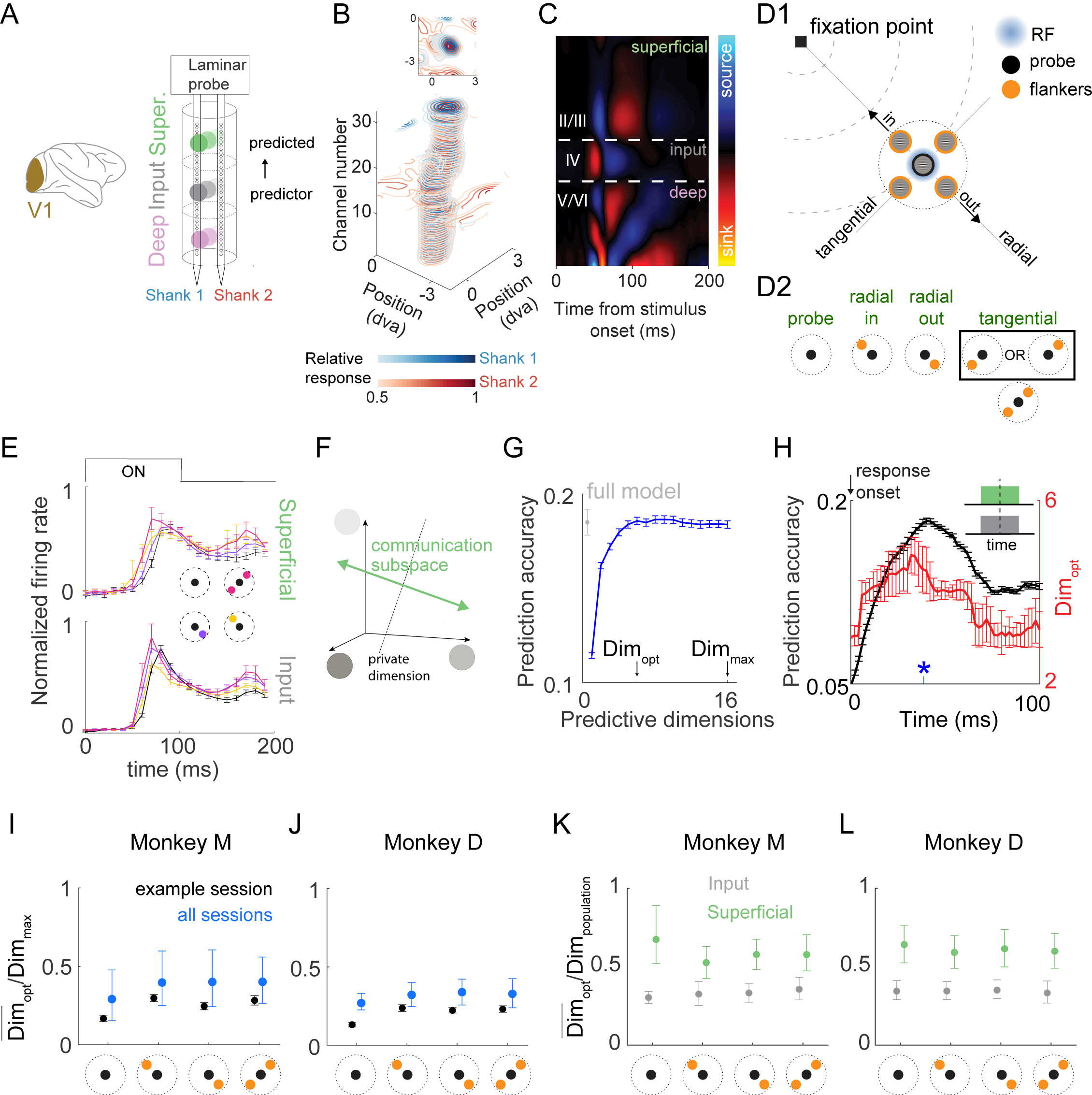
Characterization of inter-laminar communication in V1. **(A)** Illustration of electrophysiological recordings. Populations of cortical area V1 units were recorded with high-density laminar probes (2 shanks with 32 channels/shank), from two monkeys. Laminar probe simultaneously recorded population activity across the depth of the cortex. (**B**) Example receptive field contours vertically stacked for each recording site along the depth of each of the two shanks on the laminar probe. For the purpose of visualization, contours of different shanks are plotted with a small vertical offset. Top: Vertical view of the stacked contours. (**C**) Example current density (CSD) map showing the early current sink (red) indicative of the input layer. (**D**) Visual stimulation protocol. D1: Schematic of visual stimulation for the passive fixation task. The probe stimulus (black) was centered on the receptive field of the recorded neuronal populations (blue circle), in isolation or with a flanker at one of the four locations (orange) equidistant around the probe stimulus. The radial axis is defined by the center of gaze (fixation point) and probe. The tangential axis is orthogonal to it at the probe location. D2: Summary of visual conditions. In subsequent results, ‘tangential’ refers to one of the two tangential locations shown in D1, indicated by a common symbol shown in this panel. (**E**) Normalized peri-stimulus time histograms (PSTH; see Methods) of recorded units in the superficial (top) and input (bottom) layers under various visual conditions. Also shown is the visual stimulation protocol for reference. (**F**) Illustration of a low-dimensional communication subspace between cortical layers. The axes represent the activity space of input layer population (represented by three units here, for example) that are used to predict the activity of superficial layer population. Perturbations in all or fewer dimensions in this space can be predictive of the activity of superficial layer population. In this example, the green line represents the dimension along which changes in input layer activity are predictive of the activity of all units in the superficial layer, and is referred to as ‘communication subspace’. Perturbation of input layer activity orthogonal to this subspace (black dotted line) will not change the predicted activity of the superficial layer, and is referred to as ‘private dimension’. (**G**) Predicting superficial layer population activity (16 units) from input layer population activity (34 units) using reduced-rank regression (RRR) with varying number of predictive dimensions (blue curve) or a full regression model (ridge regression; grey circle) for an example session (Monkey D; under the radial-in condition). For the RRR model, *Dim_max_* is the maximal number of dimensions of source population activity patterns that can be used for prediction, and is the minimum between the numbers of the superficial and input layer units. The optimal dimensionality (*Dim_opt_*) was defined as the lowest number of predictive dimensions for which prediction performance was within one SEM of peak performance. Error bars indicate the standard error across multiple draws of trials and the corresponding cross-validation folds. (**H**) Temporal evolution of the optimal dimensionality (red) and its prediction accuracy (black) computed by RRR for the example session and visual condition used in (G). The analysis was conducted on a moment-by-moment basis using a sliding window of 50ms within the estimated responsive period (see Methods). Blue asterisk: time around which the window analyzed in (G) was centered. Error bars indicate the standard error across multiple draws of trials. (**I**) The ratio between *Dim_opt_* and *Dim_max_* under various visual conditions for an example session (black) or across all sessions (blue) from monkey M. Error bars indicate the 95% confidence interval for the mean ratio averaged across the responsive period. (**J**) Same as (I) for results from monkey D. (**K, L**) The ratio between *Dim_opt_* and the dimensionality of the population activity (*Dim_population_*) in the superficial layer (green) or the input layer (gray) across all sessions from each monkey.

Given the rich temporal dynamics exhibited by the recorded units (Fig. 1E), we hypothesized that the dynamics of inter-laminar interactions were also time-variant and therefore conducted analysis on a moment-by-moment basis. Neuronal activity was measured as spike counts in 50ms bins during the appropriate stimulus processing range of V1 neurons, which we refer to as the responsive period (see Methods). Leveraging trial-to-trial response variabilities to repeated stimuli at each spatial configuration, we characterized the inter-laminar interaction by assessing the extent to which the fluctuation of neuronal activities in the superficial layer could be predicted by those in the input layer. Specifically, we subtracted the appropriate peri-stimulus time histogram (PSTH) from each single-trial response and z-scored the residual for each type of visual stimuli separately (see Methods).

### Cortical layers interact through a communication subspace

Neuronal populations between cortical areas have been shown to interact through a ‘communication subspace’ (41, 43). That is, only a low-dimensional subspace of the upstream area neural population activity space is related to the downstream area activity (illustrated in Fig. 1F). Moreover, the dimensionality of the communication subspace from V1 to V2 has been shown to be consistently lower than the dimensionality of the target population activity (41). Based on these results, we hypothesized that cortical layers also interact through subspaces and that this is not simply due to the low dimensionality of either the source or target population activity. Specifically, we tested whether the inter-laminar interaction between the input and superficial layers in V1 was limited to a subspace of the neural activity space of the input layer by employing reduced-rank regression (RRR) (44, 45), a multivariate linear regression model with a constraint enforcing a small number of latent predictive factors (see Methods). For an example session under the radial-in condition (Fig. 1G), only 6 dimensions (*Dim_opt_*) were needed to achieve prediction performance that is as good as a full linear regression model (ridge regression; see Methods), which was lower than the maximal possible prediction dimensionality (*Dim_max_*) determined by the minimum between the number of units in the source and target populations. Moment-by-moment analysis for this session revealed that this result held throughout the responsive period of V1 (Fig. 1H). Furthermore, this result was consistent across visual conditions, sessions, and monkeys (Fig. 1I, J), implying that inter-laminar interactions in V1 shared the low-dimensional property exhibited by interareal interactions. To ensure a fair comparison across visual conditions, populations in the input and the superficial layers used to conduct prediction analysis were fixed, and the sample sizes were matched across visual conditions. These protocols prevented the number of analyzed neurons and trials from differentially affecting the analysis result (41, 46). We next tested whether this signature of inter-laminar interaction was due to a low complexity of population activity either in the source population (the input layer) or in the target population (the superficial layer). We used factor analysis to assess the complexity of population activity in either layer (see Methods). The analysis revealed that the dimensionality of activity in either population was consistently higher than the number of predictive dimensions (Fig. 1K, L). Thus, the low dimensionality of inter-laminar interactions cannot be explained by the complexity of population activity in the input or superficial layer, but rather reflected the nature of inter-laminar communications.

We next investigated the impact of the spatial configuration of visual stimuli on the structure and efficacy of inter-laminar subspace communication. Potential, non-mutually exclusive mechanisms by which inter-laminar interactions can be modulated by the spatial configuration of stimuli include: (a) the structure of the interaction is changed, which would be observable as the degraded prediction performance under a given visual condition when the corresponding input layer data was projected onto the communication subspace identified from a different visual condition, (b) the fidelity of the interaction is changed, which would be observable as differential prediction accuracies across visual conditions. To test these possibilities, we performed RRR to characterize the communication subspace and computed its prediction accuracy on a moment-by-moment basis for each visual condition.

### Structure of communication subspace is preserved across flanker locations

Testing whether the structure of inter-laminar interactions is changed requires characterizing the difference between communication subspaces identified from different visual conditions. We first investigated the relative alignment between the subspaces across different visual conditions by using the measure of principal angle (Fig. 2A), which computes angles between sequentially aligned pairs of basis vectors, each within one of the subspaces. The smallest principal angle is referred to as the ‘leading principal angle’. Small principal angles indicate a similar orientation of subspaces and imply that much of the structure of the communication subspace is preserved across visual conditions. By performing RRR using data from a sliding time window of 50ms, we identified the communication subspace for each visual condition and computed the principal angles between all possible pairs. To assess whether the obtained principal angles were significantly small, we compared them to the principal angles between randomly generated subspaces while preserving the dimensionalities of the computed communication subspaces (see Methods). As shown in Fig. 2B, pooling across all sessions, the leading principal angles between the communication subspaces identified from the probe and any type of flanked condition were consistently below chance level. The result held for all other visual condition comparisons (Fig. S1A).

**Figure 2.**
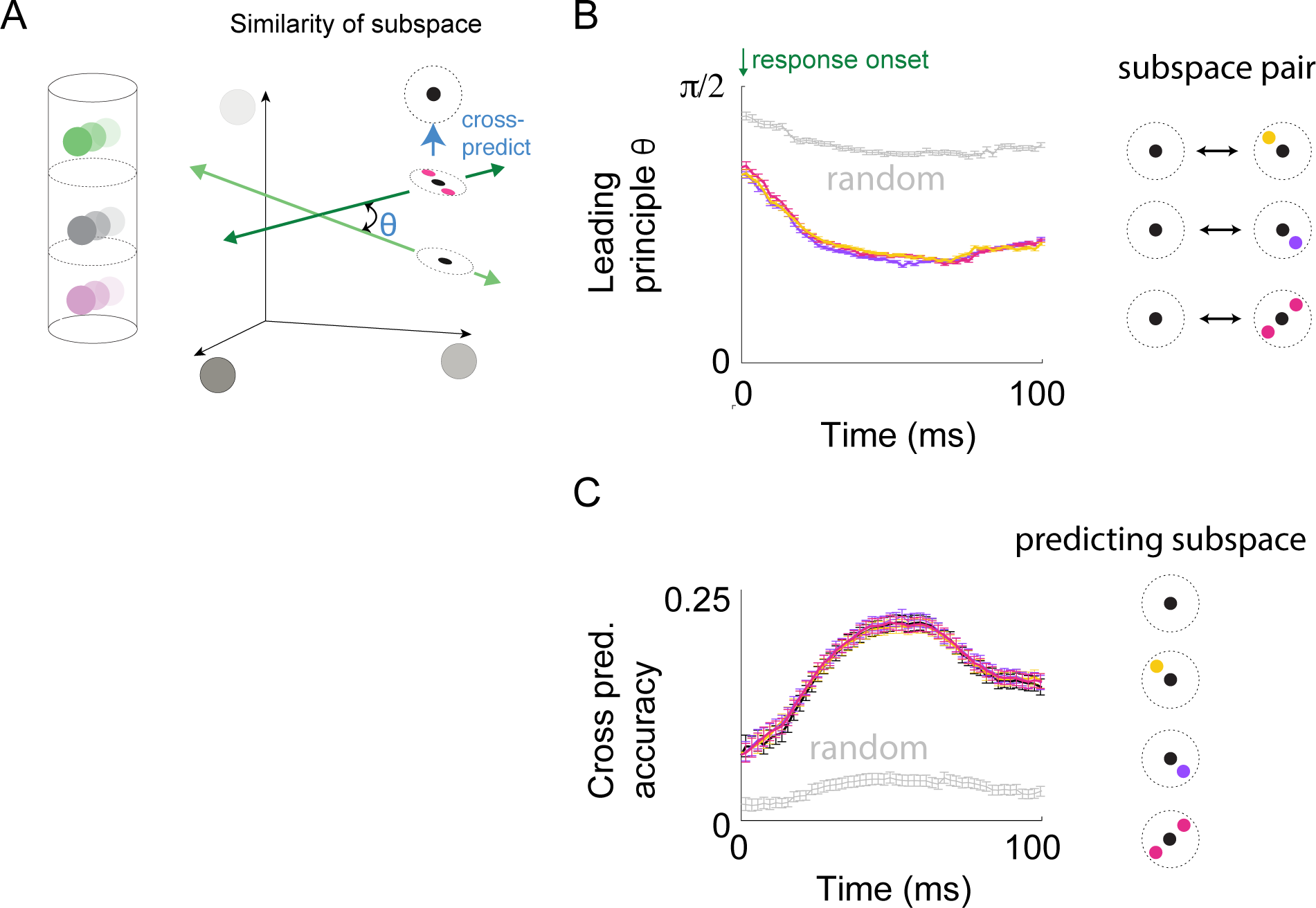
Similarity of inter-laminar communication subspace across visual conditions. **(A)** Illustration of principal angle measure and cross-prediction analysis for comparing two subspaces. Each axis represents the activity of an input layer neuron used to predict superficial layer activity. Green lines represent input-to-superficial communication subspaces identified from two visual conditions. Principal angle measures the alignment between the two subspaces. For cross-prediction analysis, input layer activity for one visual condition is first projected onto the communication subspace identified from a different visual condition, before being used to predict superficial layer activity using linear regression. (**B**) Temporal evolution of leading principal angle between the communication subspaces identified for all visual conditions (averaged across all sessions). Chance level alignment between subspaces (grey) was estimated as the leading principal angle between random subspaces of comparable ranks (see Methods). (**C**) Cross-prediction analysis for neural activity in probe condition, using projection subspaces identified from one of the four visual conditions. Error bars indicate the 95% confidence interval for the mean. Difference between conditions was inferred using estimation statistics framework (see Methods). Black: within-condition prediction, equivalent to the result obtained from RRR by definition. Yellow, purple, magenta: across-condition prediction. Grey: chance level (see Methods). For other visual condition comparisons, see Fig S1.

To characterize the influence of the relative alignment between communication subspaces across visual conditions on the strength of interaction, we performed cross-prediction analysis using the projection of the data from the source (input layer) population for a given visual condition onto the communication subspaces identified across different visual conditions using a multivariate linear regression model (Fig. 2A). Throughout the responsive period, the projected data gave similar prediction performance across all visual conditions from which the communication subspace was identified, and these were significantly above what would be expected by chance (Fig. 2C, S1B). Thus, the extent to which the input layer activity was informative of the superficial layer activity was similar among the communication subspaces identified from different visual conditions. Taken together, these two results indicate that the structure of inter-laminar interactions in a linear framework is preserved across different spatial configurations of visual stimuli.

### Efficacy of communication subspace is degraded in the presence of flankers

We next investigated how the spatial configuration of visual stimuli influenced inter-laminar prediction accuracies, starting by comparing the probe condition against the flanked condition (data pooled across all possible locations of flankers). Temporal dynamics of prediction accuracy were obtained as the predictive performance at optimal dimensionality under RRR employed on a moment-by-moment basis. As shown in Fig. 3A for a representative session, under either visual condition, the inter-laminar prediction accuracy initially increased and then decayed during the responsive period. To quantify the difference in the prediction accuracies across visual conditions for each session, we introduced a prediction modulation index (PMI; see Methods). For the example session in Fig. 3A, the corresponding PMI was significantly negative during the entire responsive period (Fig. 3B), indicating that the prediction accuracy was weakened in the presence of flankers. Despite an inter-subject difference in the initial temporal profile of the PMI, the degradation of inter-laminar prediction accuracy was consistent across sessions for both monkeys (Fig. 3C).

**Figure 3.**
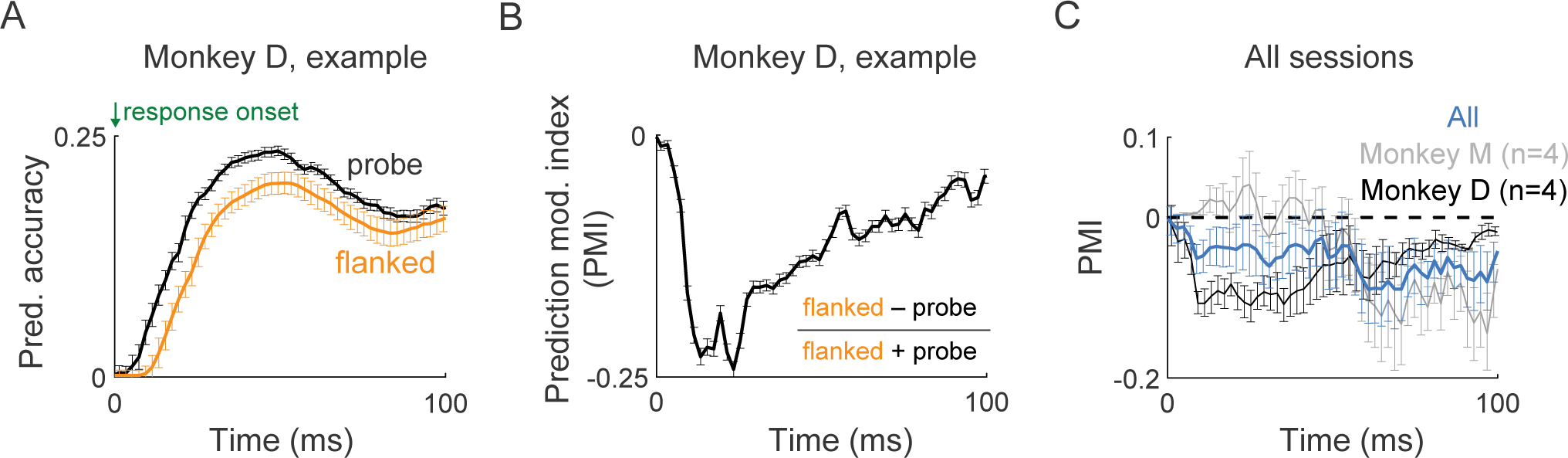
Efficacy of inter-laminar communication subspace across visual conditions. **(A)** The temporal evolution of input-superficial prediction accuracies under the probe (black) and the flanked conditions (orange) for an example session from monkey D. Error bars indicate the standard error across multiple draws of trials. (**B**) The temporal evolution of the prediction modulation index (PMI; see Methods) comparing the inter-laminar prediction accuracies under the probe and the flanked conditions. Negative PMIs correspond to a degradation of prediction accuracy under the flanked condition compared to the probe condition. Error bars indicate 95% confidence interval for the mean. Difference between conditions was inferred using estimation statistics framework (see Methods). (**C**) Same as (B) for results across sessions from each monkey (grey: monkey M; black: monkey D) or across the two monkeys (blue).

### Degradation of communication efficacy is mediated by layer-specific signals

We tested two non-mutually exclusive hypotheses about how the presence of flankers might weaken inter-laminar prediction accuracy (schematized in Fig. 4A): hypothesis *I*, the flanked condition causes the activation of a novel signal targeting the input layer, or hypothesis *II*, the flanked condition causes the activation of a novel signal targeting the superficial layer. To disentangle these possibilities, on a moment-by-moment basis, we performed the RRR analysis using data from the two layers at different temporal delays (Fig. 4B1) and thereby determined the temporal evolution of PMI as a function of delay. It is important to note here that delays associated with both inter-laminar signal conduction and intra-laminar recurrent processing would factor into this temporal analysis. If hypothesis *I* were true, the timing of the degradation and hence the temporal profile of the PMI would be independent of the inter-laminar delay being considered (Fig. 4B2, left). In contrast, if hypothesis *II* were true, i.e. if the novel signal targeted the superficial layer, the timing of the degradation would depend on the offset of the superficial layer data analyzed and therefore shift earlier with increasing temporal delay (Fig. 4B2, right). Based on these observations, we estimated the time of degradation of prediction accuracy by determining when the PMI started to decrease and related it with the temporal delay being considered. As shown in Fig. 4C for two representative sessions from the two monkeys, the decrease of PMI followed by persistently negative components arose earlier with increasing temporal delay. Across sessions, the correlation between the estimated arrival time of the novel signal and the temporal delay was consistently negative and close to -1 for both monkeys, indicating a dependence as predicted by hypothesis *II*. This result suggests that the degradation of prediction accuracy in the presence of flankers was mainly due to a novel signal targeting the superficial layer.

**Figure 4.**
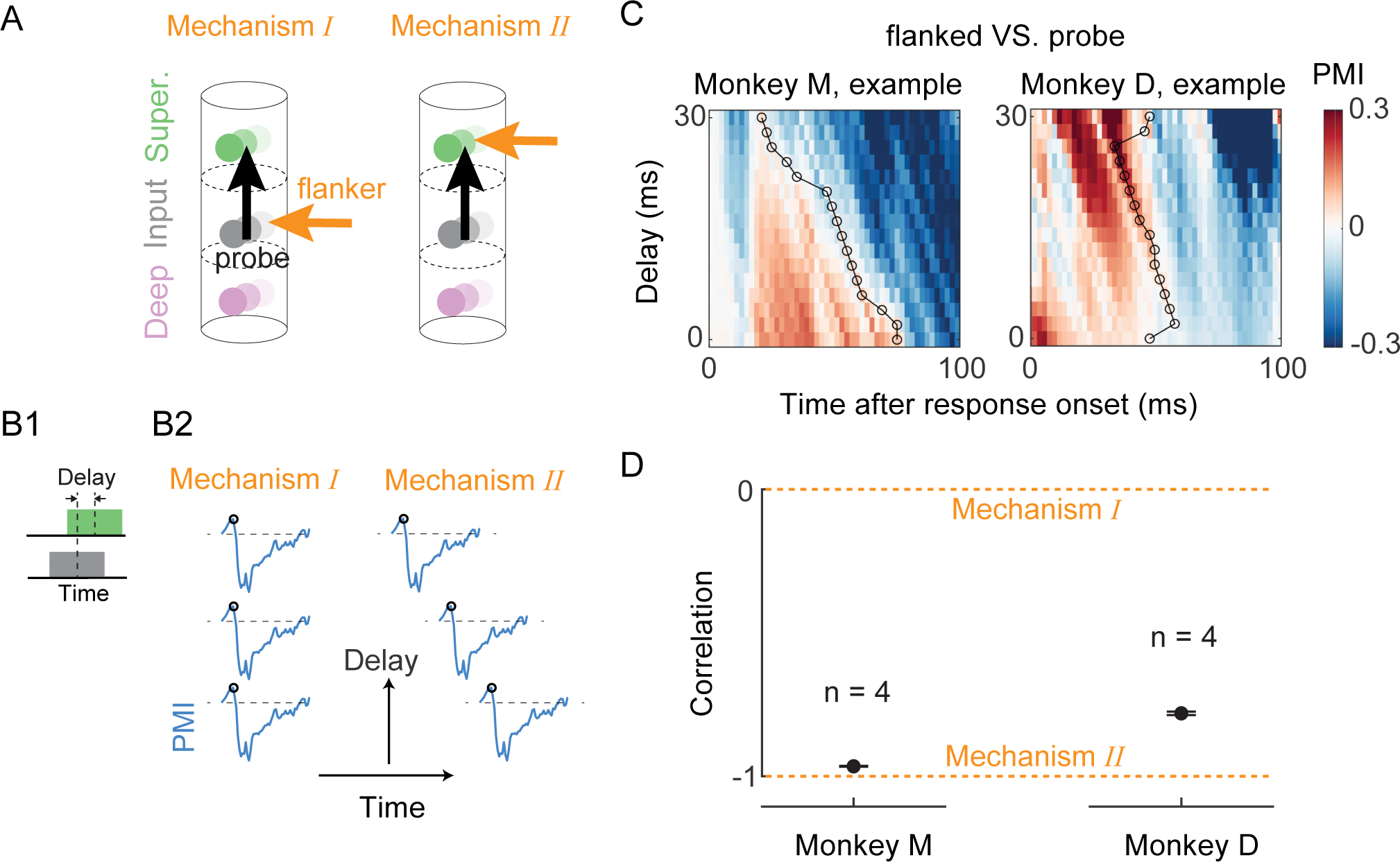
Mechanism of degradation in communication efficacy. **(A)** Illustration of two possible mechanisms by which inter-laminar communication is weakened under the flanked condition. Mechanism *I*: flanker causes the activation of a novel signal (orange arrow) targeting the input layer. Mechanism *II*: flanker causes the activation of a novel signal (orange arrow) targeting the superficial layer. (**B**) Delay analysis protocol and potential outcomes. B1: RRR was conducted on a moment-by-moment basis using input and superficial layer activity at different temporal delays. B2: Illustration of delay-induced changes in prediction accuracy modulation index (PMI) dynamics, as implied by mechanisms *I* and *II*. The onset of decreasing trend in PMI (black circle) should be independent of temporal delay under mechanism *I*, or occurring earlier with increasing temporal delay under mechanism *II*. (**C**) Temporal evolution of inter-laminar PMI under the probe and the flanked conditions at various inter-laminar temporal delays for example sessions from two monkeys. PMIs that are neither significantly positive nor negative are colored in white (the corresponding 95% confidence interval for the mean PMI includes zero). Black circle: time beyond which PMI starts decreasing and eventually becomes persistently negative at each level of delay. (**D**) Correlation between inter-laminar temporal delay and the time when PMI starts decreasing (marked in black circles in (C) for example sessions), across all sessions from each monkey. A zero correlation implies mechanism *I* while a -1 correlation implies mechanism *II*. Error bars indicate 95% confidence interval for the mean.

### Degradation of communication efficacy is sensitive to flanker location

Two robust characteristics of visual crowding identified by psychophysical studies are the asymmetry and anisotropy of crowding zones, the spatial extent over which flankers affect target recognition. First, a flanker more eccentric than the target stimulus (radial-out condition) has a greater perceptual crowding effect than an equally spaced inward (less eccentric) flanker (radial-in condition) (5). Second, the crowding zone is elongated along the radial axis so that radially positioned flankers produce a stronger crowding effect than tangential ones (tangential condition) (4). We reasoned that these perceptual asymmetries could be due to asymmetries in inter-laminar prediction accuracy in V1. In particular, we compared the prediction accuracy under the flanked condition associated with the strongest perceptual crowding effects (the radial-out condition) against the other flanked conditions (the radial-in and the tangential conditions). To ensure a fair comparison across visual conditions, we additionally aligned the temporal prediction accuracy data to account for different response latencies across conditions. We found that the prediction accuracy degraded in the radial-out condition compared to the radial-in (Fig. 5A) and the tangential conditions (Fig. 5D) over most of the responsive period. Despite an inter-subject variability in the temporal profile of the prediction degradation caused by the radial-out condition relative to the radial-in condition, such degradation was consistent across all sessions for both monkeys. Moreover, applying the inter-laminar temporal delay analysis as above, we found that the prediction degradation with radial-out flankers emerged earlier with increasing temporal delay (Fig. 5B-C, E-F), consistent with a hypothesis of a superficial-layer targeting signal that is dominant in the radial-out condition.

**Figure 5.**
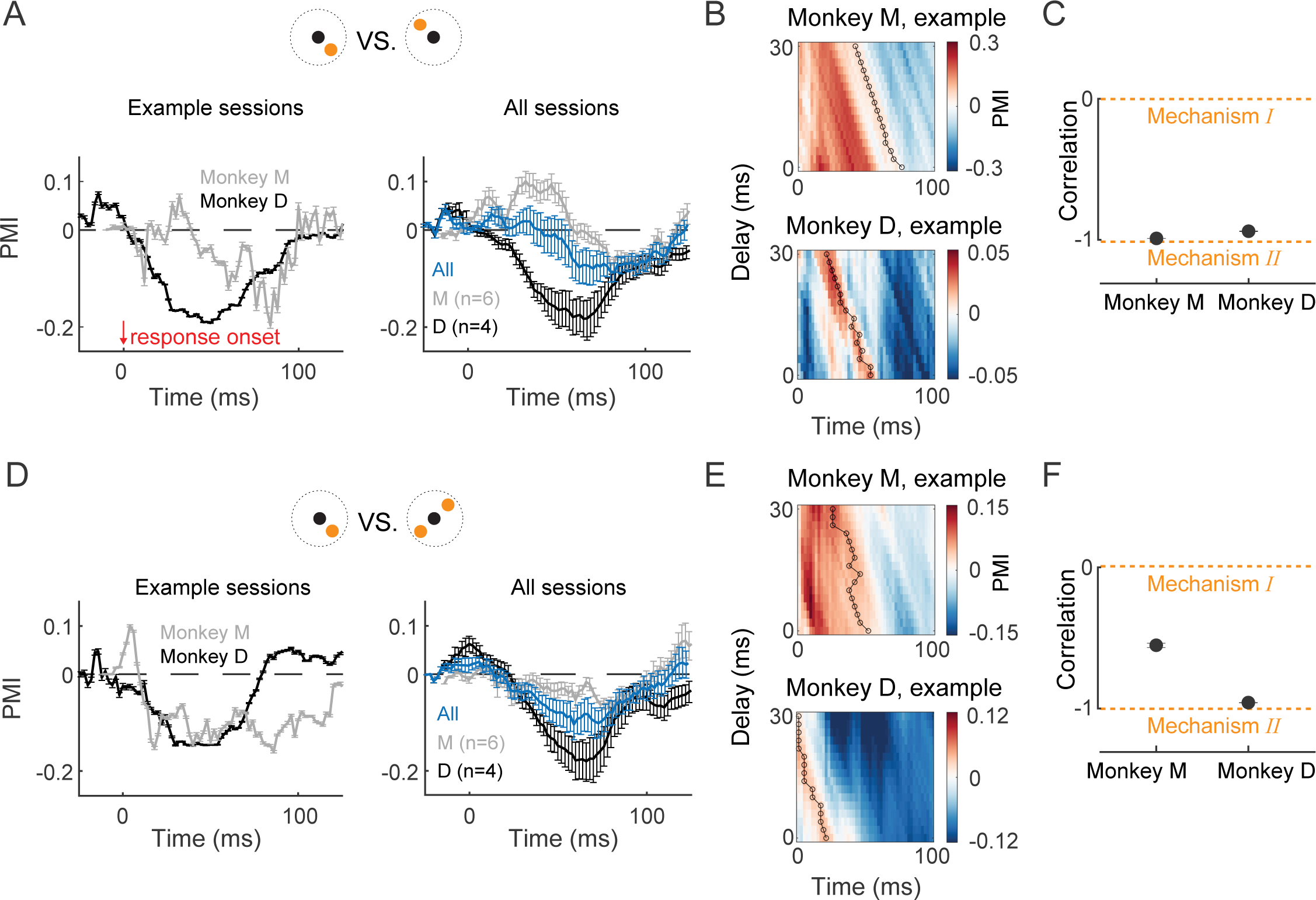
Communication efficacy degradation as a function of flanker location. **(A)** Temporal evolution of RRR prediction accuracy modulation index (PMI) comparing the radial-out and radial-in conditions for example sessions (left) or across all sessions (right), from monkeys M (grey), D (black) or both (blue). Negative PMIs imply a degradation of prediction accuracy in the radial-out condition compared to the radial-in condition. Error bars indicate 95% confidence interval for the mean. (**B**) Example sessions showing temporal evolution of PMI as a function of inter-laminar delay (see Fig. 4B, C) for the visual conditions compared in (A). (**C**) Correlation between inter-laminar delay and the time when PMI starts decreasing (marked in black circles in (B) for example sessions), across all sessions from each monkey. (**D**-**F**) Same as (A-C) comparing the modulation of inter-laminar prediction accuracy under the radial-out and tangential conditions. For comparison between the radial-in and tangential conditions, see Fig.S2A.

### Feedforward-feedback signaling balance depends on flanker location

The interplay of feedforward and feedback signaling is a hallmark of cortical information processing (47–49). Such interplay is not only prominent at the interareal level but also implicated in the layer-specific local circuit of V1 (39, 50, 51). We next sought to understand how the balance of inter-laminar feedforward and feedback signaling was affected by the spatial configuration of visual stimuli. We investigated this by employing Canonical Correlation Analysis (CCA; see Methods) to relate the activities in the input and superficial layers at different time delays on a moment-by-moment basis (Fig. 6A), which we refer to as population correlation. This methodology has been previously applied (52) to study the structure of interactions between cortical areas, finding that the balance was feedforward-dominated shortly following stimulus onset and then became feedback-dominated.

**Figure 6.**
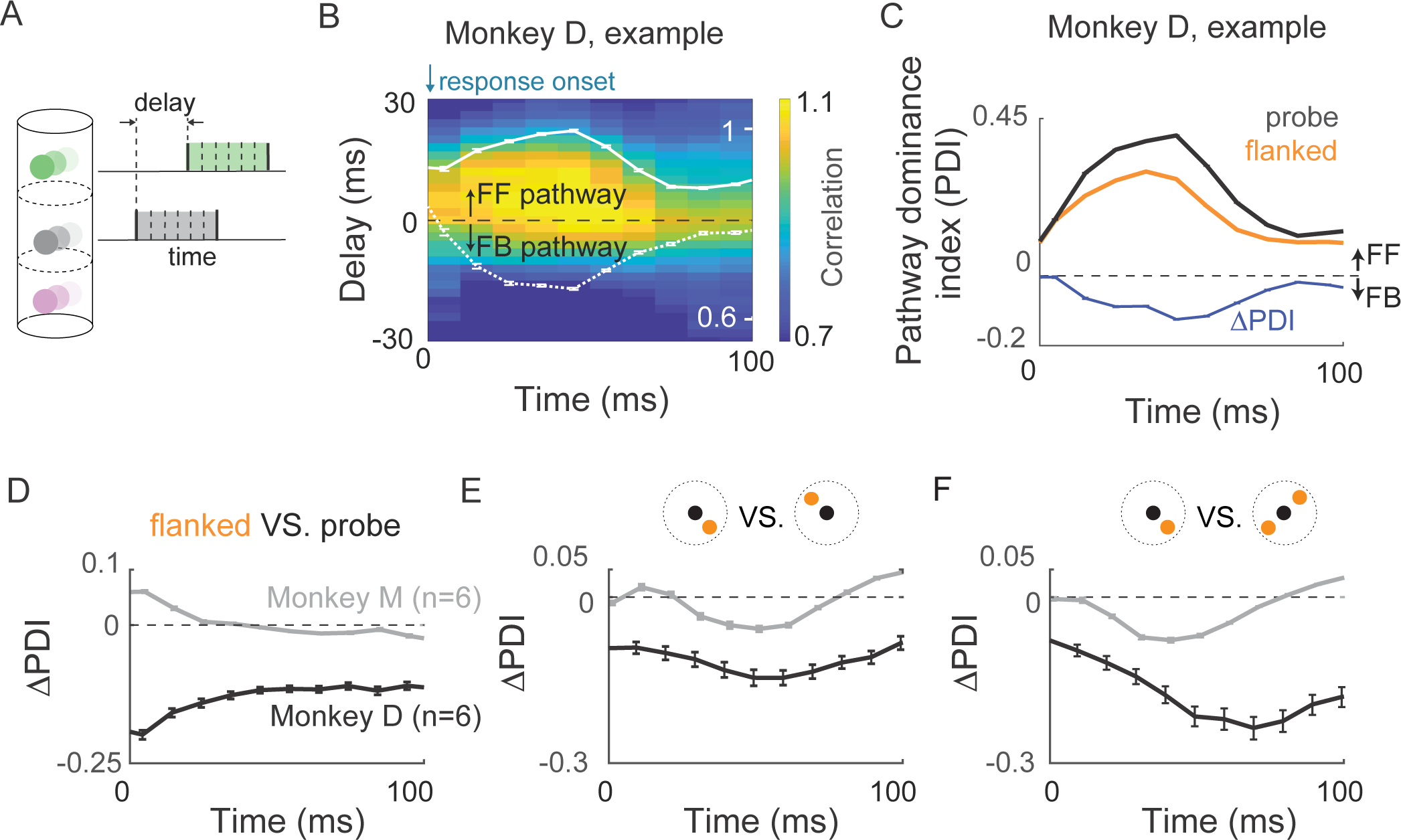
Inter-laminar feedforward/feedback balance and its modulation in the presence of flankers. **(A)** Canonical correlation analysis (CCA) protocol for estimating temporal evolution of population correlations between activities of input and superficial layers at varying temporal delays. (**B**) CCA-based population correlation as a function of time and inter-laminar delay during the visually responsive period. Overlaid traces (white) show the correlation along the feedforward (solid line) and the feedback (dashed line) directions, computed as the average correlation at positive and negative sides of delay, respectively. Error bars indicate the standard error across multiple draws of trials. (**C**) Temporal evolution of the pathway dominance index (PDI; see Methods) under the probe and the flanked conditions for an example session. A positive PDI indicates a feedforward-dominated (FF) interaction while a negative PDI indicates a feedback-dominated (FB) interaction. Also shown is the difference between PDIs across conditions (blue trace), showing the shift in balance between feedforward and feedback signaling in the presence of flankers. (**D**) Temporal evolution of the difference in the PDIs under the probe and the flanked conditions for all sessions from each monkey. (**E, F**) Same as (D) for results comparing PDIs separately under different types of flanked stimuli (**E**: radial-out VS. radial-in; **F**: radial-out VS. tangential). In (C-F), error bars indicate 95% confidence interval for the mean.

For each visual condition, we calculated the population correlation between activities in the two layers as a function of time and time delay between layers (Fig. 6B). To quantify the strength of interaction in each direction, we computed a ‘feedforward correlation’ by taking the mean over correlations for all positive delays (input layer leading superficial layer; Method) and similarly for ‘feedback correlation’ for negative delays (superficial layer leading input layer). For a representative example session under the probe condition (Fig. 6B), while the feedforward correlation increased steadily from the time of response onset and then gradually decayed, the feedback correlation was consistently lower than the feedforward correlation, indicating a feedforward-dominant interaction throughout the responsive period.

We quantified the degree to which the inter-laminar interaction was dominant in either direction of signaling by defining a pathway dominance index (PDI; see Methods), where a positive PDI indicates feedforward-dominated signaling and a negative PDI indicates feedback-dominated signaling. The larger the magnitude of PDI, the more biased the interaction was toward one direction. To investigate the impact of stimulus configuration on the balance between the inter-laminar feedforward and feedback pathways, we determined the temporal evolution of PDI separately for each visual condition. In the same example session as above, under both the probe and the flanked conditions, the PDI increased steadily and gradually decayed, remaining significantly positive throughout the responsive period (black and orange curves in Fig. 6C). Thus, the interaction became more feedforward-dominated in the early phase of the responsive period and then returned to a more balanced interaction. Remarkably, the PDI was less positive under the flanked condition, indicating that the balance was shifted away from the feedforward direction in the presence of a flanker. This result was robust across sessions for each monkey (Fig. 6D). Furthermore, similar analysis revealed that this modulation was flanker position-dependent. In the visual condition associated with the strongest perceptual crowding effect (the radial-out condition), the balance was shifted away from the feedforward direction compared to both the radial-in and tangential conditions (Fig. 6E, F). Thus, the non-uniformity in stimuli-specific prediction accuracy analysis (Fig. 5) was mirrored by the modulation of the interplay between inter-laminar feedforward and feedback signaling. Interestingly, the inter-subject difference in the temporal profile of the difference in PDI across visual conditions (probe VS. flanked, radial-out VS. radial-in) was similar to that in PMI obtained from the prediction analysis, such that both the shift of balance (Fig. 6D and Fig. 6E) and the degradation of inter-laminar prediction accuracy (Fig. 3C and Fig. 5A) emerged later during the responsive period for Monkey M compared to Monkey D.

### Strength of charge sink in superficial layer is sensitive to flanker location

To test the hypothesis that our observations are a reflection of a signal that targets the superficial layers and is sensitive to flanker locations, we next examined the relative strength of this signal evoked by flanker-only stimuli at different locations. For each flanker-only condition, we determined the level of charge sinks by integrating over time the early current sinks in the superficial layer obtained from CSD response (Fig. 7A-C; see Methods), which reflected the subthreshold integrated input to local neurons (42, 53). For both monkeys, flankers at the radial-out position evoked stronger charge sinks compared to flankers at the radial-in or tangential positions (Fig. 7D), thus providing direct evidence for a location-dependent superficial-layer targeting signal that differentially impacts the representation of the target. These results support a hypothesis that the observed degradation in communication efficacy and shift in feedforward-feedback signaling balance are mediated by retinotopically non-uniform cortical connectivity in the output layers of V1 (Fig. 7E).

**Figure 7.**
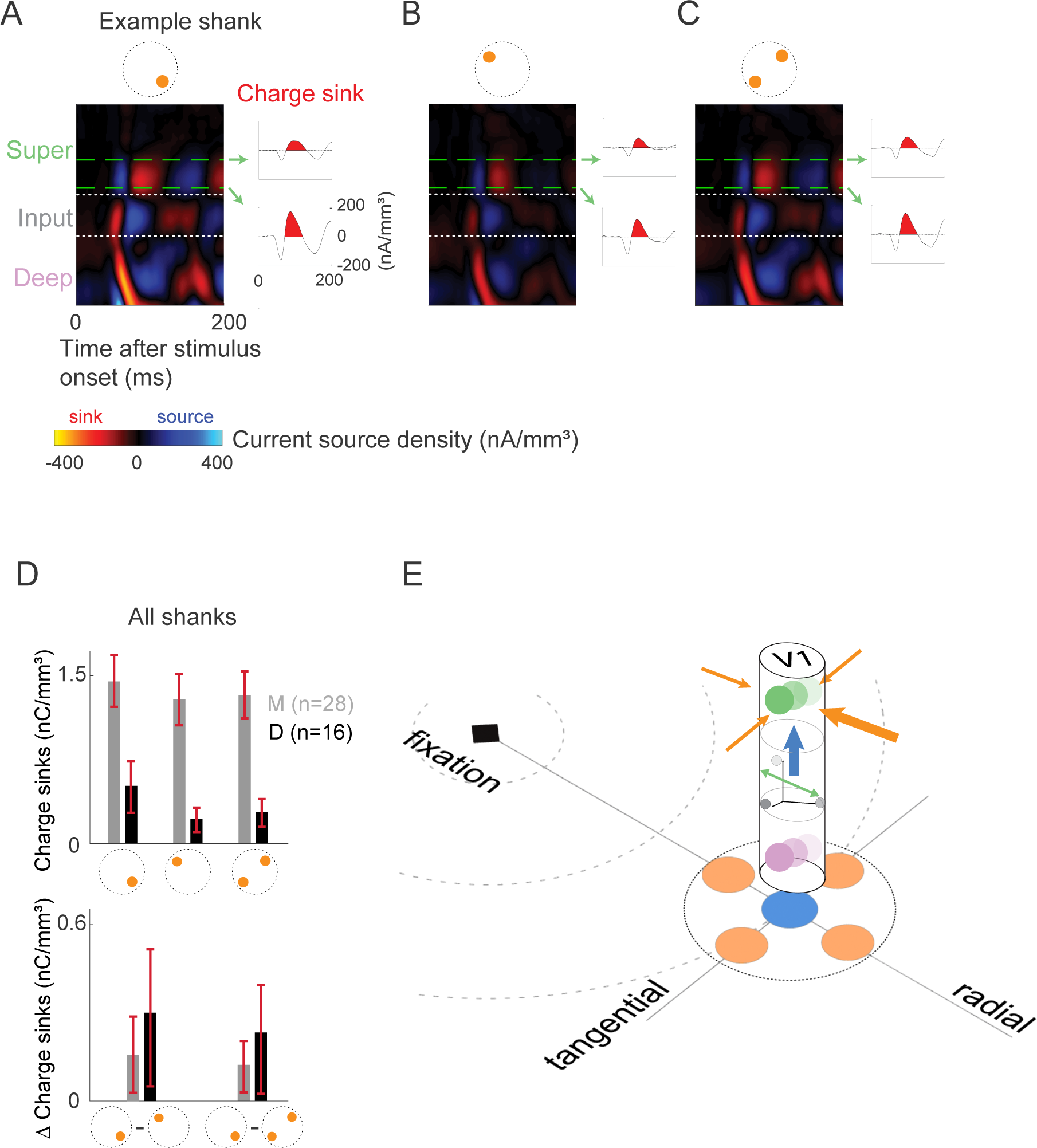
Retinotopically non-uniform context integration in V1. **(A)** An example of estimated CSD across V1 layers evoked by flanker presented by itself at the radial-out position. Dotted white lines indicate laminar boundaries (see Methods). Traces on the right show temporal evolution of the CSD signal at two recording sites on the example shank, marked by the green dashed lines. The earliest charge sink (area shaded in red) reflects integrated subthreshold input to neuronal ensembles as a direct consequence of visual stimulation. (**B, C**) Same as (A) with the flanker presented at the radial-in and tangential positions, respectively. (**D**) Top: Level of early charge sinks under different flanker-only conditions across all sessions from each monkey. Bottom: Within-shank difference in the level of early charge sinks between the radial-out and radial-in conditions (left), the radial-out and tangential conditions (right). Error bars indicate 95% confidence interval for the mean. Difference between conditions was inferred using estimation statistics framework (see Methods). For comparison between the radial-in and tangential conditions, see Fig.S2B. (**E**) Summary of findings. The inter-laminar interaction between the input and superficial layers in V1 is limited to a subspace of the neural activity of the input layer, whose structure does not change in the presence of contextual stimuli. Instead, contextual stimuli modulate its efficacy through the non-uniform activation of novel signals targeting the superficial layer.

## DISCUSSION

We leveraged simultaneous laminar recordings to understand how the spatial configuration of visual contextual stimuli affected inter-laminar interactions in area V1 of the macaque. V1 activity has been extensively studied as a locus of surround modulation (33–40) and more recently has been implicated as a bottleneck impairing perception under visual crowding (6–9, 13). We established that V1 laminar populations interact through a communication subspace. We demonstrated that inter-laminar communication efficacy was degraded in the presence of contextual stimuli. This degradation was not accompanied by changes in the structure of the subspace of neural activity along which the inter-laminar interaction occurs. Furthermore, we found that the balance between the inter-laminar feedforward and feedback signaling was non-uniformly shifted in the presence of flankers. Strikingly, these modulations matched the spatially non-uniform aspects of perceptual degradation, such that a greater degree of modulation was associated with a flanker at the visual location that is known to exert a stronger perceptual impairment. Finally, we found that the spatial configuration of contextual stimuli differentially modulated contextual signaling in the superficial layers. Our results suggest a model in which degraded communication in the sensory hierarchy, mediated by retinotopically non-uniform connectivity in the output layers of V1, underlies the perceptual impairments in spatial vision.

### Contextual modulation of information flow

Despite decades of research at the level of perception, investigation into the neural mechanisms of non-uniform perceptual degradation in peripheral vision has received limited attention. The most studied hypothesis is that the perceptual degradation is mediated by changes in the tuning properties of neurons and therefore leads to information loss about target features as early as the input and superficial layers of V1 (6, 7, 54). The observed impairment of information encoding has been shown to be greater with flankers positioned at visual configurations that exert stronger crowding effects, either with shorter target-flanker distance or at a radial-out location relative to the target stimulus. However, modulation of neural responses and changes in information coding do not necessarily imply changes in signaling fidelity along processing stages in the visual hierarchy. Our results demonstrate that inter-laminar interactions in V1, a key mechanism of hierarchical signaling, are disrupted by spatial context, which may account for the accumulation of information loss along the visual hierarchy (7).

Although we address degraded inter-laminar interaction in V1, our results do not rule out the possibility of additional mechanisms responsible for visual crowding in higher visual areas. A stronger information loss in area V4 has been observed with crowding due to the summation of signals within the larger receptive fields of V4 neurons compared to V1 (7). Other relevant studies relying on coarser measures of neural activity such as fMRI, found that interareal temporal correlations (between V1, V2, V3, V4 and the visual word form area) are lower with crowded letters compared to uncrowded letters (14). Such deterioration in the extrastriate cortex can only compound the degradation of signaling we identified within V1, as higher visual areas ultimately rely on V1 inputs for their computations (1).

### Source of non-uniform context signal

The dependence of the PMI temporal profile on the inter-laminar delay suggested that the degradation of prediction accuracy in the presence of flankers was mainly due to a novel signal targeting the superficial layer. This result is broadly consistent with a recent study (36) that examined the laminar profile of current sinks in the CSD upon stimulation of the receptive field surround with isotropic annular gratings and found, based on onset latency measurements, that the processing of such spatial context initiates in the superficial and deep layers. Our results significantly extend this prior work by (a) characterizing the effect of such a mechanism on information propagation along the intra-V1 hierarchy, and (b) showing that such a mechanism could also be a template for non-uniform contextual modulation. Further experiments are needed to be able to pinpoint the source of this implicated input, which could non-exclusively be horizontal connections from superficial layer neurons outside the recorded V1 column or feedback connections from higher visual areas. Both possibilities are supported by previous studies in the context of surround modulation where optogenetic inactivation of horizontal connections in mouse L2/3 V1 (55) or marmoset V2 feedback connections to V1 (50) reduced the amplitude of surround modulation. Moreover, each type of connection was shown to contribute to the processing of spatial context at different spatiotemporal scales (36, 40). It is important to note that our results do not conflict with previous work suggesting the contribution of geniculate feedforward connections, which primarily terminate in the input layer of V1, to the processing of contextual stimuli (33, 39, 56–59), but imply a weaker effect of such connections on input-superficial interactions compared to connections terminating in the superficial layers. Interestingly, our observation of an intermediate level of negative correlation (neither close to 0 nor to -1) between the time when degradation emerged and the temporal delay being considered for the comparison between the tangential and radial-out conditions from one monkey (Fig. 5F), suggests the influence of potentially both types of connections on the anisotropy. Thus, the mechanism underlying the non-uniform aspect of the modulation of inter-laminar interaction by contextual stimuli could vary with the specific locations of flankers being compared.

### Features of inter-laminar communication

Our study has identified two signatures of inter-laminar interactions in V1: low-dimensionality and feedback signaling. By employing RRR, we demonstrated that the interaction between the input and superficial layers occurred through a low-dimensional communication subspace, akin to inter-areal interactions (41, 43) but in contrast to interactions within the superficial layer of V1 (41). The low-dimensional structure could confer the computational benefit of flexible and selective routing of activity to downstream targets.

Extensive studies have attempted to infer feedback interactions between brain areas by relating activities between areas with temporal delay (52, 60–64), computing phase delays in local field potentials (LFP) or multiunit neuronal activity (MUA) (51, 65–67), and comparing the timing of neuronal response onsets (68–70) as well as the emergence of certain neuronal response properties (71–74) across areas. Yet feedback interactions among within-area laminar circuits in general, and between the input and superficial layers in particular, remain unknown. Here we characterized inter-laminar interactions along both feedforward and feedback directions by applying CCA with varying temporal delay between layers. Notably, we observed substantial levels of correlation over a range of negative delays, especially at the initial and late phases of the responsive period, implying a feedback component (superficial leading input) in the interlaminar interaction. This result is consistent with the implication of a previous study that examined the laminar profile of the multiunit neuronal activity (MUA) for the alpha rhythm and found that MUA in the superficial and deep layers preceded MUA in the input layer (51). Given that the dendritic arbors of the input layer (L4C) neurons are locally confined and the descending axons of the superficial layer neurons mainly pass through the input layer with very weak branching (28, 29, 75), the identified feedback signaling is less likely to be relayed via a direct superficial-input anatomical connection. Instead, certain types of neurons in the deep layer, the primary target of projections from neurons in the superficial layer, have been shown to send extensive axonal projections into the input layer (28, 76, 77), which form an indirect superficial-input pathway via the deep layer and could serve as an anatomical substrate for the identified feedback signaling.

In conclusion, this study provides compelling evidence that contextual interactions in the visual cortex exhibit anisotropy at the earliest stages of the visual hierarchy. Flankers modulate the input-superficial inter-laminar interaction in a spatially anisotropic manner. Radial-out flankers most effectively weaken the fidelity and shift the directionality of the interaction while preserving its structure. Contextual signaling in the superficial layers is retinotopically non-uniform. These findings advance our understanding of the neural bases of spatial vision and offer mechanistic insights into the processing of contextual information in the visual cortex. More broadly, they shed new light on the retinotopic organization of the visual cortex.

## AUTHOR CONTRIBUTIONS

XX, ASN & MPJ conceptualized the project. MPM & ASN designed the experiments. MPM and NVH collected data and performed preliminary analyses. XX conducted the statistical analysis in the study. ASN & MPJ supervised the experimental and analytical aspects of the project, respectively. XX, ASN & MPJ wrote the manuscript.

## ACKNOWLEDGEMENTS

This research was supported by NIH/NEI R01 EY032555, NARSAD Young Investigator Grant, Ziegler Foundation Grant and Yale Orthwein Scholar Funds to ASN, NIH/NEI R01 EY034605 and R00 EY025026 to MPJ, the Kavli Institute for Neuroscience at Yale University (Kavli Postdoctoral Fellowship to XX), the Swartz Foundation (Swartz Postdoctoral Fellowship in Theoretical Neuroscience to XX), NIH/NINDS training grants T32-NS007224 and T32-NS041228 to MPM, and by an NIH/NEI core grant for vision research P30 EY026878 to Yale University. We would like to thank the veterinary and husbandry staff at Yale for excellent animal care.

## INCLUSION AND ETHICS

We support inclusive, diverse and equitable conduct of research.

## MATERIALS & METHODS

### Data Collection

#### Surgical procedures

Surgical procedures were similar to those described previously (78–80). We placed low-profile titanium recording chambers in two rhesus macaques so that the chambers allowed access to area V1 (both left and right hemispheres in monkey M, right hemisphere in monkey D). Chambers were targeted based on sulcus reconstructions created using preoperative structural MRI. After chamber implantation, we removed the native dura mater and replaced it with a transparent silicone artificial dura (AD). The AD allowed for the visualization of cortical sites in V1 for probe targeting. All procedures were approved by the Yale University Institutional Animal Care and Use Committee and conformed to NIH guidelines.

#### Electrophysiology

Prior to a series of recordings, we electroplated (nanoZ, White Matter LLC) 64-channel electrode arrays (“laminar probes,” NeuroNexus Technologies, Inc., 2 shanks, 32 channels/shank, 70µm spacing between sites, 200µm between shanks) with PEDOT (poly(3,4-ethylene dioxythiophene)). At the beginning of each recording session, we inserted a laminar probe in V1. Laminar probes were attached to a titanium mounting stage that was screwed into the chamber. We positioned the laminar probes using an electronic micromanipulator (Narishige Inc.) and ensured that the probes were orthogonal to the surface of the cortex by visual inspection through a surgical microscope (Leica Microsystems). To position the probe within the brain, we first penetrated the AD, arachnoid, and pia by moving the probe downward at a high speed (>100µm/s). After the tip of the probe entered the cortex, we inserted the remainder of the probe at a slow speed (2µm/s). Once the entire probe was in the brain, we slowly (2µm/s) relieved the pressure on the brain by retracting the probe upward, relieving pressure on the brain but not moving the probe relative to the cortex.

Electrical signals from the laminar probe were collected at 30kHz and digitized on a 64-channel digital headstage and sent to the recording system (RHD Recording System, Intan Technologies). Action potential waveforms were extracted offline using Kilosort2 (81, 82) and manually sorted into single and multi-unit clusters (Phy; 81, 82). Clusters with peaks preceding the trough were identified as axonal spikes and excluded. Recordings were collected over the course of 22 sessions (14 in monkey M, 8 in monkey D) with hundreds of units recorded in each subject (693 single units and 430 multi-unit clusters in monkey M, 400 single units and 277 multi-unit clusters in monkey D).

#### Behavioral Control and Eye Tracking

We controlled behavioral experiments using NIMH MonkeyLogic (83). Eye position and pupil diameter were sampled at 120Hz (ETL-200, ISCAN Inc.) and sent to the behavioral control system. Stimuli were presented on a monitor positioned 57cm from the monkey with a 60Hz refresh rate. Trials were aborted if the eye position deviated more than 1.2-1.5 degrees of visual angle (dva; 1.2 for monkey D, 1.5 for monkey M) from the central fixation point.

#### Receptive Field Mapping

We mapped RFs of the column by presenting Gabor patch stimuli (2-4 cycles/degree, 0.25-1 degree Gaussian half-width, 100% luminance contrast) on a square grid spanning the lower visual quadrant of interest (both left and right in monkey M, left in monkey D) while the monkey maintained fixation on the center of the screen. Grid spacing parameters were optimized each session and ranged from 0.25-1 dva. A stimulus was presented at a random location and orientation on the grid during each frame. We calculated the LFP power for each recording channel 40-200ms after stimulus presentation in each location. The LFP power at each location was smoothed using a Gaussian kernel (sigma = 0.75 dva), and the peak location averaged across all recording sites was defined as the receptive field center. Spatial receptive field maps for each channel were plotted as stacked contours for each shank for visualization.

#### Current Source Density Mapping

We used CSD mapping (42) to identify laminar boundaries in our recordings and estimated the relative strength of the signal targeting the superficial layer across visual conditions.

While monkeys maintained fixation on the screen, 100% luminance contrast white annular stimuli were flashed for 32ms, positioned at the center of the RF. The LFP signal following the stimulus onset was averaged across trials and spatially smoothed using a Gaussian kernel (sigma = 140µm). The CSD was calculated as the second spatial derivative of the LFP:

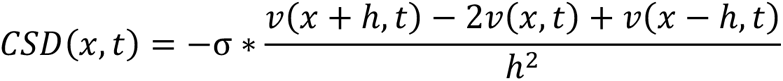

where *x* is the position in the extracelluar medium at which CSD is calculated, *t* the time following the stimulus onset (advancing in 1 ms), *h* the spacing between recording sites on the linear probe (here 70 μm), *ν* the voltage, σ the conductivity of the cortical tissue (0.4 S/m). We interpolated the CSD every 7 μm. The input layer was identified by the boundaries of the early current sink characterizing feed-forward input into layer IV. Channels above and below this sink were classified as superficial and deep, respectively. CSD provides a link between LFP and neuronal ensemble activities by approximating the net local transmembrane currents which generate the local LFP. The identified current sinks (negative deflections, visualized in red in Fig. 7A-C) in the extracellular medium reflect integrated subthreshold input to neuronal ensembles (42, 53). We used CSD response to flanker-only stimuli (radial-out, radial-in, tangential) to estimate the relative strength of the signal targeting the superficial layer evoked by contextual stimuli at different locations. In particular, for each shank on the laminar probe, we first identified the early current sink (within 200ms following stimulus onset) in its CSD response. At each depth along the superficial layer, we calculated the level of charge sinks by integrating the current sink over time and then averaged the result across the depth of the superficial layer. Since the distance between flanker and the receptive field of the recorded V1 column here is smaller compared to a previous study of monkey V1 where a surrounding stimulus presented alone in the absence of receptive field stimulation did not evoke significant spiking activity from the recorded cells (36), in some sessions, we still observed the earliest sinks in the input layer. Therefore, the identified CSD sinks in the superficial layer may involve feedforward input from the input layer, which we assumed was independent of flanker location. Comparing the level of charge sinks evoked by flankers at different locations relative to the receptive field, thus provides information about the relative strength of contextual signal targeting the superficial layer.

#### Experimental Task

While monkeys maintained fixation, stimulus arrays were flashed on the screen for 100 ms and off the screen for 200-250 ms. The arrays consisted of a target stimulus in the receptive field center either in isolation (probe condition) or together with a flanking stimulus (flanked condition). There were four different conditions (probe, tangentially flanked, radially inward flanked, and radially outward flanked) of stimulus arrays presented during a trial. This was repeated 4-6 times during each trial depending on the monkey’s ability to hold fixation. The stimulus conditions were randomly interleaved from flash to flash. The center of the target stimulus was aligned with the center of the average of the RFs of each recording site. The four flanker locations were positioned on the radial or tangential axes. The radial axis was defined as the axis connecting the target (and RF) center and the fixation point at the center of the monitor. The tangential axis was defined as the line orthogonal to the radial axis and passing through the target center. The tangential flanked condition occurred when the target was presented along with a flanker on the tangential axis, either in the clockwise or counterclockwise direction. The radial-in flanked condition occurred when the target was presented along with a flanker on the radial axis and between the target center and the fixation point. The radial-out condition was similar to the radial-in condition, except the flanker was placed on the radial axis further from the fixation point than the target. Target-flanker spacing was identical across stimulus conditions.

The probe condition was presented on 10% of flashes. On 80% of flashes, flanked stimuli were presented. In the flanked condition, the flanker was positioned at a tangential, radial-in, or radial-out location with equal probability. If the flanker was presented tangential to the target, it was positioned clockwise or counterclockwise to the target with equal probability. In the remaining 10% of trials, a stimulus was presented exclusively at one of the four flanker locations. Target stimuli were sine Gabor patches (25% luminance contrast, 3.5 cycles/degree, 0.5-1.0 degree Gaussian half-width) presented at 6 evenly spaced orientations and 2 opposing phases. Flankers were presented at 100% luminance contrast but were otherwise identical to the target stimuli. There was a 0.1-0.2 degree gap between the edges of the target and flankers (edge defined as 2 standard deviations from the center of the Gaussian). In the flanked condition, flankers were presented at the same orientation as the target or orthogonal to the target with equal probability. When flankers were presented alone, they were shown with the same orientation distribution as the targets.

### Data Analysis

Wherever possible, instead of relying on null-hypothesis testing, we show bootstrapped estimations of differences between conditions using estimate statistics framework (84, 85), which provides a principled way to measure effect sizes coupled with estimates of uncertainty, yielding interval estimates of uncertainty. Since the experiment assigned substantially different probability of being presented for the probe (10%) and each type of flanked condition (26.7%), when comparing inter-laminar interactions for a pair of visual conditions, we drew the same number of trials under each condition multiple times and conducted the analysis with 20-fold cross validation.

#### Normalized peri-stimulus time histograms (PSTH)

PSTHs were generated based on spike counts in 30ms bins shifted by 10ms. The PSTH for each neuron was calculated separately for each visual condition and normalized between the maximal PSTH over time under the probe condition and the baseline firing rate taken as the PSTH from -30ms to 30ms relative to stimulus onset. Neurons whose PSTHs never surpassed the 95% confidence interval of their baseline firing rates were excluded. The averaged normalized PSTH exhibits a maximum less than 1 because of the variability in the peak time across neurons.

#### Response onset estimation

For each recorded unit, we estimated its response onset under each type of visual condition. We took the PSTH from -30ms to 30ms relative to stimulus onset as the baseline firing rate. The response onset was identified as the time when the PSTH surpassed the 95% confidence interval of the baseline firing rate. Units whose PSTH never crossed the threshold were taken as non-responsive units. For each session, the response onset of the simultaneously recorded population in a specific layer was estimated by taking the average of response onsets of all responsive units. An interval of 100 ms following the response onset of the input layer population was considered as the stimulus processing range (referred to as ‘responsive period’) of V1 for the purpose of characterizing the inter-laminar interaction between the input and superficial layers.

#### Unit selection for regression analysis

For comparison of the inter-laminar prediction accuracy across visual conditions, we only included those units in the superficial layer well predicted by the recorded population in the input layer, which were identified by employing a Lasso regression with 20-fold cross-validation to each recorded unit in the superficial layer within the estimated responsive period. The prediction performance was calculated under the largest *λ* value such that the normalized squared error (*NSE*) was within one standard error of the minimum *NSE* across *λ* parameters. Units in the superficial layer with normalized squared error smaller than 1 were identified as being well-predicted.

#### Data preparation for regression analysis

We counted spikes in a sliding window of 50ms within the estimated responsive period. We investigated how the neuronal activities in the input and superficial layers were related by assessing the extent to which trial-to-trial fluctuations of population responses in the superficial layer could be predicted by that in the input layer of V1. Therefore, for each type of visual stimuli (orientation, spatial configuration, orientation difference between target stimulus and flanker), we subtracted the appropriate peri-stimulus time histogram (i.e., average time-varying signal; PSTH) from each single-trial response and z-scored the residual. For all analyses, we excluded units with low firing rates (less than 0.2 spikes/s on average).

#### Regression

To assess the extent to which trial-to-trial fluctuations of population responses in the superficial layer could be predicted by that in the input layer of V1, we first applied a linear model of the form *Y = XB* using ridge regression, which was referred to as the full regression model. To test whether the inter-laminar interaction occurred through a ‘communication subspace’, we used reduced-rank regression (RRR), which constrains the linear weight matrix B to be of a given rank. The details of this analysis can be found in (41).

Both ridge regression and RRR were applied to a sliding window of 50ms (advancing in 2ms) in both layers with 20-fold cross validation. The prediction accuracy was calculated as 1 − *NSE*, where *NSE* is the mean normalized squared error between the test data and the predictions across folds. For RRR, we took the smallest number of dimensions for which predictive performance was within one SEM of the peak performance as the optimal dimensionality. To study the effect of the spatial configuration of visual stimuli on prediction accuracy, we analyzed the temporal evolution of the prediction accuracies at optimal dimensionality across visual conditions. For each pair of visual conditions, we first aligned the temporal evolutions of corresponding prediction accuracies to compensate for the difference in the response onsets across visual conditions and computed a prediction modulation index, which was calculated as:

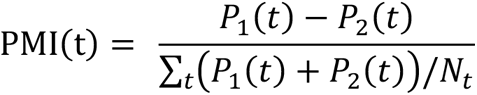

Where P_1_(t) and P_2_(t) are the prediction accuracies at optimal dimensionalities obtained by RRR at time *t* after response onset under visual conditions 1 and 2, respectively. *N_t_* is the number of timepoints involved in the analysis.

#### Factor analysis

We quantified the complexity of neural activity in the superficial layer of V1 by using factor analysis (FA), a dimensionality reduction technique which allows spiking variability to vary across neurons and calculates the dimensionality of covariance. The details of this analysis can be found in (46, 86, 87). We followed the same steps as previously published work (41, 43) to estimate the dimensionality: We first determined the number of dimensions *m_peak_* that maximized the cross-validated log-likelihood of the observed residuals. Then we fitted an FA model with *m_peak_* dimensions and chose *m*, by the eigenvalue decomposition, as the smallest dimensionality that captured 95% of the variance in the shared covariance matrix. These population dimensions (*m*) and predictive dimensions as determined from RRR are based on different techniques.

#### Principal angle

We characterized the relative alignment of two communication subspaces identified from different visual conditions by using the measure of principal angle, which computes angles between sequentially aligned pairs of basis vectors, each within one of the subspaces, so as to minimize the angle between them. Notably, this measure does not require the compared subspaces to have the same rank. The smallest angle obtained was taken as the ‘leading principal angle’. The details of this method can be found in (88, 89). A small leading principal angle implies similar orientations of the communication subspaces identified from different conditions. To verify that the experimentally obtained leading principal angles were significantly small, we compared them to the principal angles between subspaces that were randomly generated but with preserved ranks of the actual communication subspaces under different visual conditions. On a moment-by-moment basis, we generated and compared for each pair of visual conditions 5000 pairs of randomly generated subspaces, from which the leading principal angle and its confidence interval (returned by estimate statistics) were calculated.

#### Cross-prediction analysis

An additional analysis to compare the communication subspaces was based on measuring the inter-laminar prediction performance under a given visual condition when the corresponding input layer data was projected onto the communication subspace identified from a different visual condition. We computed the prediction performance for when projecting the input layer data from the communication subspace identified from condition A onto that identified from condition B (across-condition prediction performance). Then we compared it to the prediction performance when the input layer data was projected onto the original communication subspace of condition A (within-condition prediction performance; obtained from RRR by definition). Comparable cross-condition and within-condition prediction performance implies a similarity between the structure of communication subspaces identified from different conditions. To verify whether the computed cross-condition and within-condition prediction performances were significantly close to each other, we compared them to the prediction performance obtained by chance, where the input layer data in a given condition was projected onto subspaces randomly generated as in the principal angle analysis. Note that this analysis was performed twice for each pair of visual conditions, considering either condition as the within-condition communication subspace.

#### Population correlation analysis

We used canonical correlation analysis (CCA) (90) to capture population correlation between the input and superficial layers at different time delays on a moment-by-moment basis. CCA finds pairs of dimensions, one each in the neuronal activity space in each layer, such that the correlation between the projected activity onto these dimensions is maximally correlated. The exact description and formulation of CCA can be found in (52). We took two windows of activity, one in each layer. Window length was 50ms and the window was advanced in 10ms steps. The activity within each window was then binned using 10ms bins. The reported results were robust over a reasonable range of window and binning length chosen. We reported correlation associated with the first two canonical pairs (correlations associated with the third were on average 60% lower and close to chance level).

The correlations along the feedforward (FF) and the feedback (FB) signaling pathways were calculated as the mean correlation at positive and negative delays, respectively:

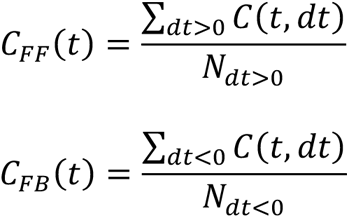

where *t* is the time following response onset, *dt* is the inter-laminar delay involved between windows from two layers, *C(t, dt)* is the corresponding correlation. *N_dt_*_>0_ is the number of positive delays investigated, which is equal to the number of negative delays, *N_dt_*_<0_

The pathway dominance index was calculated as:

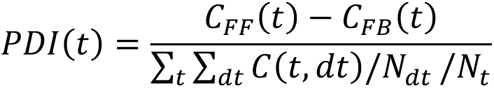

where *N_t_* and *N_dt_* are the number of timepoints and delay involved in the analysis, respectively. A positive *PDI* indicates a feedforward-dominated signaling and a negative *PDI* indicates a feedback-dominated signaling.

---

**Figure S1.**
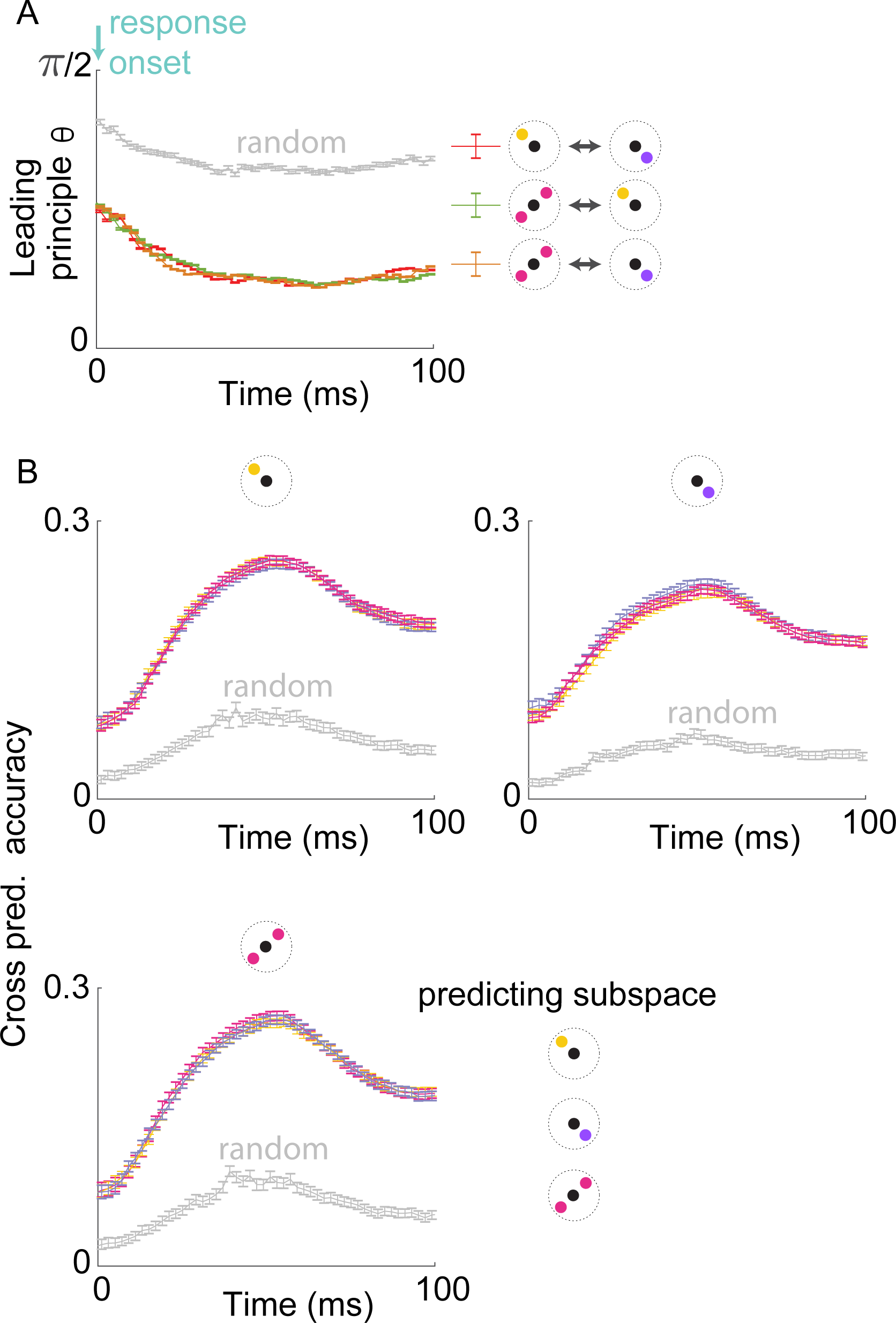
Comparing communication subspace across visual conditions. **(A)** The temporal evolution of leading principal angles between the communication subspaces identified from different types of flanked conditions during the corresponding visually responsive period (averaged across all sessions) and between random subspaces of comparable dimensions (grey; see Methods). (**B**) Cross-prediction analysis for input layer predicting superficial layer activity during a given type of flanked condition (shown at the top) and using subspaces identified for various conditions (as shown in the legend). Error bars indicate the 95% confidence interval for the mean.

**Figure S2.**
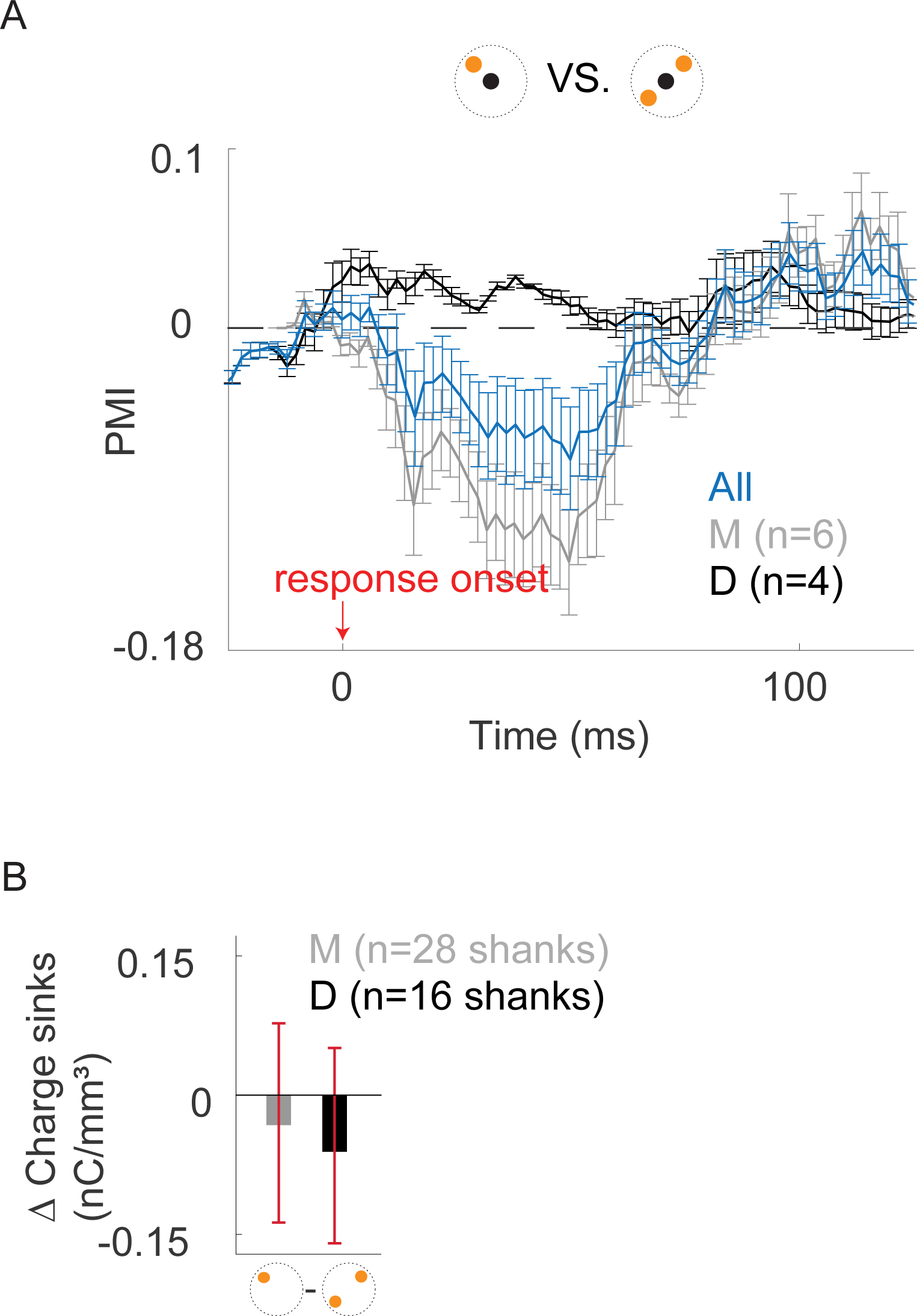
Comparing prediction accuracy across visual conditions. **(A)** Same as Fig. 5A, comparing inter-laminar prediction accuracy under the radial-in condition to that under the tangential condition. Negative PMIs imply a degradation of prediction accuracy in the radial-in condition compared to the tangential condition. (**B**) The within-shank difference in the level of early charge sinks in the superficial layer, when compared between the radial-in and tangential conditions for all shanks from each monkey. Error bars indicate 95% confidence interval for the mean.

